# Characteristics of fMRI responses to visual stimulation in anesthetized vs. awake mice

**DOI:** 10.1101/2020.08.17.253500

**Authors:** Thi Ngoc Anh Dinh, Won Beom Jung, Hyun-Ji Shim, Seong-Gi Kim

## Abstract

The functional characteristics of the mouse visual system have not previously been well explored using fMRI. In this research, we examined 9.4 T BOLD fMRI responses to visual stimuli of varying pulse durations (1 – 50 ms) and temporal frequencies (1 – 10 Hz) under ketamine and xylazine anesthesia, and compared fMRI responses of anesthetized and awake mice. Under anesthesia, significant positive BOLD responses were detected bilaterally in the major structures of the visual pathways, including the dorsal lateral geniculate nuclei, superior colliculus, lateral posterior nucleus of thalamus, primary visual area, and higher-order visual area. BOLD responses increased slightly with pulse duration, were maximal at 3 – 5 Hz stimulation, and significantly decreased at 10 Hz, which were all consistent with previous neurophysiological findings. When the mice were awake, the BOLD fMRI response was faster in all active regions and stronger in the subcortical areas compared with the anesthesia condition. In the V1, the BOLD response was biphasic for 5 Hz stimulation and negative for 10 Hz stimulation under wakefulness, whereas prolonged positive BOLD responses were observed at both frequencies under anesthesia. Unexpected activation was detected in the extrastriate postrhinal area and non-visual subiculum complex under anesthesia, but not under wakefulness. Widespread positive BOLD activity under anesthesia likely results from the disinhibition and sensitization of excitatory neurons induced by ketamine. Overall, fMRI can be a viable tool for mapping brain-wide functional networks.

## 1. Introduction

Understanding how vision works is one of the most fundamental and important objectives in the field of neuroscience (Huberman & Niell, 2011; Seabrook, Burbridge, Crair, & Huberman, 2017). Because mouse models are extensively used for biomedical research due to their cost effectiveness and the abundance of available genetic/molecular tools, the mouse visual system has been extensively investigated using conventional neuroscience tools, including electrophysiology (Grubb & Thompson, 2003; Niell & Stryker, 2008; L. P. Wang, Sarnaik, Rangarajan, Liu, & Cang, 2010). Mice have two visual pathways from the retina to the primary visual cortex (V1): the direct cortical-geniculate and the indirect colliculo-cortical pathways (Beltramo & Scanziani, 2019; Huberman & Niell, 2011; Seabrook et al., 2017). The cortical-geniculate pathway passes through the dorsal lateral geniculate nucleus (LGd), which relays retinal information directly to the visual cortex, but it also receives feedback information from the cortex and other subcortical areas, such as the superior colliculus (SCs) (Grubb & Thompson, 2003; Seabrook et al., 2017). The colliculo-cortical pathway passes through the SCs, and it indirectly relays information to cortical areas, including the primary visual and higher-order visual areas, via the lateral posterior nucleus (LP) and LGd (Seabrook et al., 2017). Eighty-five to ninety percent of mouse retinal input is sent directly to the SCs. These pathways can be mapped noninvasively and repeatedly using functional magnetic resonance imaging (fMRI) to provide useful insights into the development and reorganization of visual functions in normal, transgenic, and diseased mouse models.

To the best of our knowledge, only a few previous fMRI studies have investigated the mouse visual system (Huang et al., 1996; Lee, Li, Coulson, & Chuang, 2019; Niranjan, Christie, Solomon, Wells, & Lythgoe, 2016). The first mouse visual fMRI study, performed under sodium pentobarbital anesthesia (Huang et al., 1996), reported a negative blood oxygenation level dependent (BOLD) response in the superior portion of the brain and positive responses in the section mostly anterior to the cerebellum. In a more recent mouse fMRI study performed under medetomidine sedation (Niranjan et al., 2016), the BOLD response in the LGd and SCs increased to 10 Hz with the stimulation frequency, whereas that of the V1 decreased with stimulation frequency and was negative at 10 Hz. However, those frequency-dependent responses differed from published electrophysiologic measurements in awake and urethane-anesthetized mice (tuning frequency: 6 – 4 Hz in the LGd and 3 Hz in the V1) (Durand et al., 2016). The major reason for this discrepancy was probably the anesthetics used for the fMRI studies because they affect neural activity and neurovascular coupling (Masamoto, Fukuda, Vazquez, & Kim, 2009; Masamoto & Kanno, 2012).

The selection of an appropriate anesthetic is critical to obtain reliable, physiologically relevant fMRI responses. Many anesthetics, including urethane, propofol, etomidate, isoflurane, and medetomidine (Bosshard et al., 2010; Nair & Duong, 2004; Petrinovic et al., 2016; Reimann & Niendorf, 2020; Schlegel, Schroeter, & Rudin, 2015; Schroeter, Schlegel, Seuwen, Grandjean, & Rudin, 2014), have been used in fMRI studies of the mouse brain during paw stimulation. However, the results of such studies have been inconsistent, and bilateral activity caused by cardiovascular responses has frequently been reported (Reimann et al., 2018) when only the contralateral hemisphere was expected to be active. Recently, we found that ketamine/xylazine anesthesia produces reliable localized somatosensory-induced fMRI responses (Jung, Shim, & Kim, 2019; Shim et al., 2018; Shim, Lee, & Kim, 2020) without changing the cardiac pulsations or respiration rate (Shim et al., 2018). Thus, we used ketamine/xylazine anesthesia for this mouse fMRI study to characterize the BOLD fMRI responses to visual stimulation. Because anesthetic agents affect neural activity and hemodynamic responses, the effect of anesthetics on fMRI should be investigated by comparing fMRI under anesthesia and wakefulness.

In the present study, we developed an awake mouse fMRI protocol to minimize head motion, characterized 9.4 T BOLD fMRI responses to visual stimuli of varying pulse durations and temporal frequencies under anesthesia, and compared those fMRI responses with responses from wakeful mice. The characteristics of mouse BOLD responses were determined and compared with previously published mouse electrophysiological data (Durand et al., 2016; Grubb & Thompson, 2003; Piscopo, El-Danaf, Huberman, & Niell, 2013). Unexpected interesting observations were: 1) anesthesia induced sustained positive BOLD responses to visual stimulation in the V1, whereas wakefulness induced biphasic or negative BOLD responses, and 2) regions supposed not to respond to flashing visual stimuli were active during visual stimulation under anesthesia, but not under wakefulness. These diffused BOLD responses under ketamine can be explained by the action of the anesthesia, similar to previously reported neurophysiological findings (Haider, Hausser, & Carandini, 2013; Vaiceliunaite, Erisken, Franzen, Katzner, & Busse, 2013).

## 2. Materials and methods

### 2.1. Animal preparation

Twenty-eight male C57BL/6 mice, eight to fifteen weeks of age (23 – 28 g; Orient Bio, Seongnam, Korea), were used with approval from the Institutional Animal Care and Use Committee of Sungkyunkwan University and in accordance with the standards for humane animal care. All experiments were performed in compliance with the guidelines of the Animal Welfare Act and the National Institutes of Health Guide for the Care and Use of Laboratory Animals.

#### Head holder implantation

Thirteen animals (seven awake and six anesthetized) were used for fMRI studies. To minimize head motion during fMRI scanning, we built a polyether ether ketone (PEEK)-based, 1.25-mm-thick head post with an 8mm-diameter center hole to minimize space between the surface coil and the brain (Fig. 1A). To secure the head post onto the dorsal skull, we adopted the standard aseptic surgical procedures reported in (Chen et al., 2020; Schlegel et al., 2018). Animals were initially anesthetized with ketamine/xylazine anesthesia (100 mg/kg and 10 mg/kg, respectively, intraperitoneal [IP]). After placing the mouse in a stereotactic frame, hair was shaved, and lidocaine hydrochloride 2% was subcutaneously injected under the scalp. All soft tissues were removed and washed away by saline with cotton buds. After ensuring that the skull was completely clean, a thin layer of etching gel (Dentin Etchant Gel, Sun Medical Co., Ltd, Japan) was applied to fill small dents in the skull and minimize captured air bubbles. Then, a thin layer of light-curing (blue light) self-etch adhesive (Adper Easy One, 3M ESPE AG, Seefeld, Germany) was applied to the exposed skull, and the head post was placed on the exposed skull. All the exposed region, including the head post, was covered with a smooth layer of dental cement to reduce distortion in the MRI images caused by air. After the surgery, the mice were allowed to recover from anesthesia and then returned to their cages. Habituation for the fMRI scans was started after seven days of recovery.

**Figure 1.**
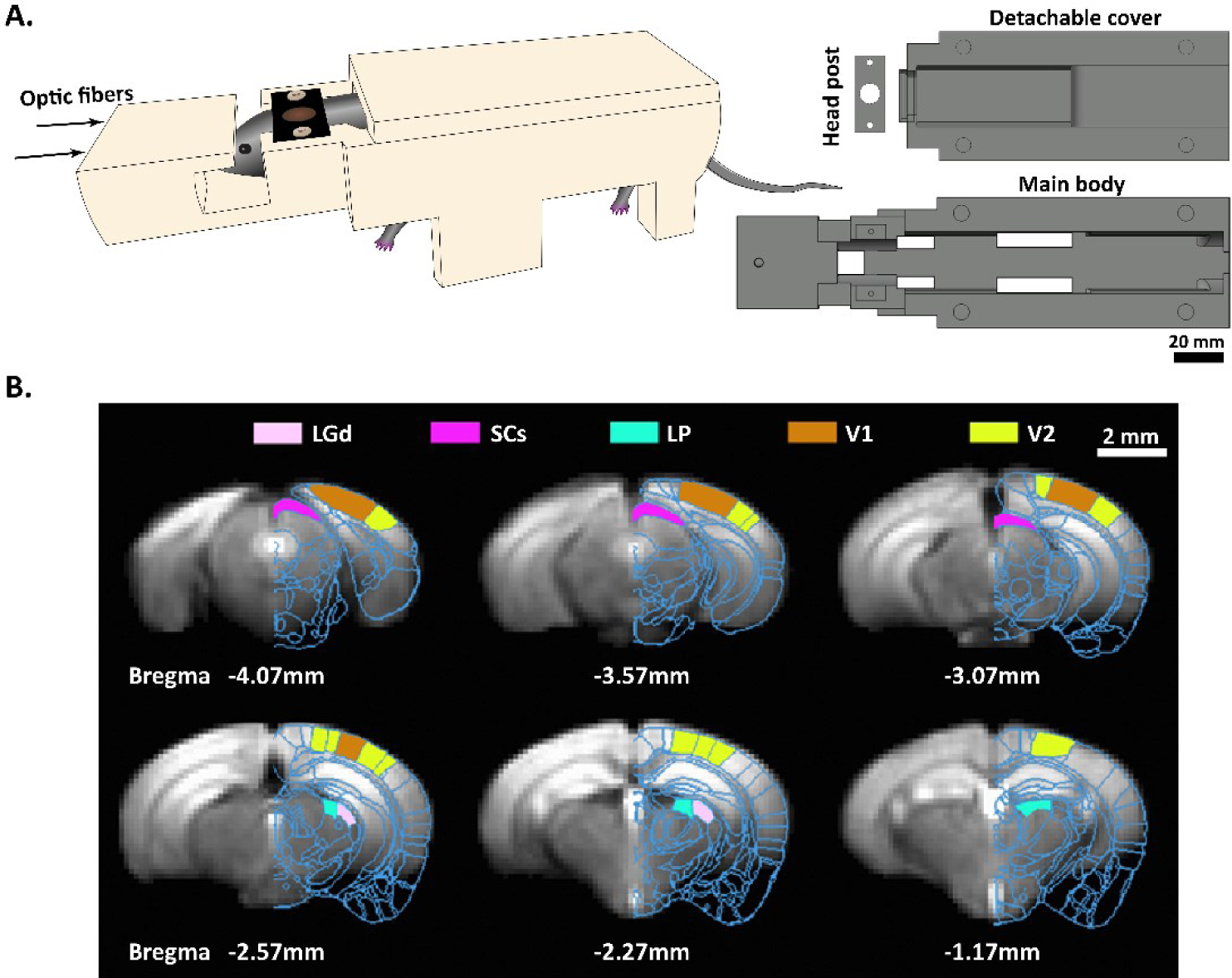
MRI-compatible cradle and definition of regions of interest in the visual areas. (A) The body restraint cradle (left) and schematics for three cradle pieces (right) specifically designed for the awake mouse fMRI in this study. The head post (black) attached to the mouse head was screwed to the body restraint cradle (ivory), which accommodated free paw movements (left). The body restraint cradle consisted of a main body and a detachable cover, and the head post had an 8-mm-diameter hole. The top view of the main body shows the four slots for positioning paws (right). (B) Regions of interest (ROIs) overlaid on an EPI template: dorsal lateral geniculate nuclei (LGd), superior colliculus (SCs), lateral posterior nucleus of the thalamus (LP), primary visual area (V1), and higher-order visual area (V2). The same ROIs in both hemispheres were used for further data analyses. The Allen Mouse Brain Atlas of a corresponding slice (Allen Institute for Brain Science, http://mouse.brain-map.org/) was overlaid on the right hemisphere.

#### Preparation and habituation protocol for awake mouse fMRI

An awake mouse cradle (Fig. 1A) that could be screwed to the head post was designed to minimize motion during the fMRI scans. The cradle consisted of two pieces: i) a main body component whose inner diameter fit to the mouse body, with open slots in the bottom for freely movable paws, and ii) a detachable cover piece to restrain body and neck motion. The same cradles were used for habituation and the fMRI studies. In all habituation and scanning sessions, the mice were initially anesthetized with 1% isoflurane then positioned in the cradle and fixed by the screw. After isoflurane anesthesia was terminated, the animals woke up before starting habituation. All habituation session were started at 1 pm every day to match circadian rhythms (Gong et al., 2015; Yoshida et al., 2016). Seven mice were habituated for ten consecutive days from an initial 30 min session to 120 min sessions, as described in Table 1. For the first seven days, each habituation period was incremented with 15 min in a mock scanner; 110 – 120 dB noises recorded during actual fMRI experiments using echo-planar imaging (EPI) sequence were introduced on day 3, and visual stimulation was additionally applied from day 7. For the three remaining days, four mice (Group 1) continued their training in the mock scanner, and three mice (Group 2) were habituated in the 9.4 T scanner to better imitate actual fMRI conditions.

**Table 1.** Daily habituation procedure for awake mouse fMRI

| Training day | 1 | 2 | 3 | 4 | 5 | 6 | 7 | 8 | 9 | 10 |
| --- | --- | --- | --- | --- | --- | --- | --- | --- | --- | --- |
| Duration (min) | 30 | 45 | 60 | 75 | 90 | 105 | 120 | 120 | 120 | 120 |
| **EPI sound** | - | - | + | + | + | + | + | + | + | + |
| **Visual stimulus** | - | - | - | - | - | - | + | + | + | + |

#### Ketamine/xylazine anesthesia protocol

For the anesthetized mouse fMRI, twenty-one mice in total were used: fifteen for typical anesthetized fMRI experiments and six for comparison of awake vs. anesthesia. The detailed anesthetized mouse fMRI protocol was reported previously (Shim et al., 2018). In short, an initial IP injection of ketamine and xylazine (100 mg/kg and 10 mg/kg, respectively) was administered. Each mouse with trimmed whiskers was placed in a customized cradle with earplugs and a bite bar. To maintain anesthesia during the scan, supplementary doses (25 mg/kg ketamine and 1.25 mg/kg xylazine) were infused intermittently (every 30 or 45 minutes) through an IP line, as deemed necessary based on changes in physiological parameters (Jung et al., 2019; Shim et al., 2018; Shim et al., 2020).

#### General experimental procedure

In all experiments, the animals were able to breathe spontaneously and were supplied with 80% air/20% oxygen at a rate of 1 L/min using a nose cone. The eyes of the mice were covered with eye gel (Hanlim Pharmaceutical Co., Ltd) to keep them moist. Respiration and temperature were continuously measured using a physiological monitoring system (Model 1030, Small Animal Instrument Inc., Stony Brook, USA) with a pressure-sensitive sensor and a rectal temperature probe, and the data were recorded using a data acquisition system (Acknowledge, Biopac Systems, Inc., Goleta, CA, USA). Body temperature was maintained at 37±0.5°C with a warm-water bath (Bruker BioSpin, Billerica, MA, USA).

### 2.2. MRI experiments

All MRI experiments were performed using a horizontal bore 9.4T/30 cm Bruker (Bruker BioSpec, Billerica, MA, USA) with an actively shielded 12-cm-diameter gradient operating with a maximum strength of 66 G/cm and a rise time of 141 μs. A combination of an 86-mm inner diameter (ID) volume coil for radio frequency (RF) transmission and a 10-mm ID surface coil for RF reception was used. The surface coil was positioned on top of the posterior side of the mouse’s head to cover the visual area.

The magnetic field homogeneity was shimmed globally using the field map method. Initially, T1-weighted sagittal images were obtained. Then, 15 contiguous coronal slices 0.5-mm thick were selected. T1-weighted anatomical images were acquired using a fast low angle shot (FLASH) sequence with field of view (FOV) = 15 × 15 mm^2^, matrix size = 256 × 256, in-plane resolution = 0.059 × 0.059 mm^2^, repetition time (TR) = 300ms, echo time (TE) = 3 ms, and number of averages (NA) = 4. Before starting the fMRI experiments with an EPI sequence, local field homogeneity was optimized in the ellipsoidal volume covering the visual area using the Bruker MAPSHIM shimming protocol (ParaVision 6, Bruker BioSpec, Billerica, MA, USA). All fMRI data were acquired using single-shot gradient-echo EPI with nine contiguous slices without gaps in the coronal plane, FOV = 15 × 10 mm^2^, acquisition matrix = 96 × 64, in-plane resolution = 0.156 × 0.156 mm^2^, slice thickness = 0.5 mm, TR/TE = 1000/20 ms, flip angle = 50°, NA = 1, sampling bandwidth = 300 kHz, and 10 dummy scans.

### 2.3. Experimental designs for visual stimulation

For visual stimulation, optic fibers with 0.5-mm diameter were placed 2 cm distally from each eye of the animal and connected to a cold white LED driver (DC200, Thorlabs, Newton, New Jersey, USA). The illuminance of the LED light was measured to be around 10 lux at the fiber tips. The LED driver was controlled by a pulse generator (Master 9; World Precision Instruments, Sarasota, FL, USA). Image acquisition and stimulus application were synchronized. Three fMRI experiments were performed, and six to eight trials were obtained for each experimental condition.

#### Experiment #1

Pulse duration dependency (n = 7 animals). To investigate the pulse duration-dependent BOLD fMRI responses, three different light pulse durations (1 ms, 10 ms, and 50 ms) were used at a 3 Hz frequency. The order of the different durations was pseudo-randomized. Each fMRI trial consisted of a 40-s pre-stimulus, 15-s stimulus, and 40-s post-stimulus period.

#### Experiment #2

Frequency dependency during anesthesia (n = 8 animals). Five stimulation frequencies (1 Hz, 3 Hz, 5 Hz, 8 Hz, and 10 Hz) delivered at a 1 ms pulse duration were used to characterize the frequency-dependent responses. The order of different frequencies was pseudo-randomized, and the same experimental design as given for *Experiment #1* was used.

#### Experiment #3

Anesthetized vs. awake condition (n = 13 animals). Seven mice were used for the awake fMRI, and six mice were anesthetized. Stimulation was delivered at 10 ms of pulse duration with two different frequencies (5 Hz and 10 Hz). Each fMRI trial consisted of 30 s (off), 10 s (on), 40 s (off), 10 s (on), and 30 s (off). The experimental time was limited to 2 hours.

### 2.4. Motion-sensitive data analyses

Pressure-sensitive signals were processed by a bandpass filter, followed by detrending. Under anesthesia, mice had no movements except the abdominal movements caused by respiration; therefore, the pressure-sensitive data were used to extract respiratory rates. Motion-sensitive respiratory rates were averaged over repeated fMRI trials in the same imaging session, producing averaged respiratory rates for each subject. In the awake animals, the pressure-sensitive signal was prone to movements caused by the freely moving paws and breathing, and thus we used them to screen large body motions.

### 2.5. Functional imaging data analyses

All data analyses were performed with the Analysis of Functional NeuroImages package (Cox, 1996), the FMRIB Software Library (Smith et al., 2004), Advanced Normalization Tools (Avants et al., 2011), and Matlab® codes (Mathworks, Natick, MA, USA).

Data underwent the following preprocessing steps to minimize noise fluctuations and improve the detection of activation: slice timing correction, image realignment, and linear detrending for signal drift removal. EPI images were registered to an EPI general space, which was made by averaging the EPI images obtained from fifteen mice. Six motion parameters (3 translations and 3 rotations) were obtained in each fMRI trial. Also, frame-wise displacements (FD) were calculated from fMRI data using the sum of the absolute displacements of the six motion parameters for each animal and assuming that the mouse brain is a sphere with a radius of 5 mm (Belloy et al., 2018; Jung et al., 2019; Power, Barnes, Snyder, Schlaggar, & Petersen, 2012). After the six motion parameter regression was applied to reduce motion artifacts (Chen et al., 2020; Satterthwaite et al., 2013), the time series were normalized by the average of the pre-stimulus baseline volumes.

In the awake studies, data screening was necessary to minimize artifacts from large head motions. If FD was ≥ 0.15 mm at any time-series points (equivalent to one voxel), then that specific trial was discarded. For the successful trials that passed the motion threshold, a mean FD value was determined across 120 time-series points for each trial, and the median of those trial-wise mean FD values was calculated. When an fMRI trial had a mean FD value close to the median of the trial-wise mean FD values, then its first volume was used as the reference volume. Afterward, all trials were realigned to the reference volume in the awake condition. In the anesthetized condition, the first volume of the first fMRI trial was used for the realignment reference volume because none of the trials had significant motion. Then, all trials with the same stimulation paradigm within an imaging session were averaged.

#### Group analysis

Animal-wise functional maps were calculated using a general linear model (GLM) analysis with a two-gamma function; the statistically significant activation threshold was set to an uncorrected p < 0.05 and cluster size > 10 voxels. The group activation maps were generated using a one-sample t-test and taking into account the significance of the family-wise error (FEW) corrected to p < 0.05. Spatial smoothing using a Gaussian kernel with 0.2 mm full-width at half-maximum (FWHM) was included to minimize potential misalignments and enhance the statistical power. For visualization, fMRI activation maps were overlaid on mouse brain EPI template images.

### 2.6. Quantitative region of interest analysis

#### Selection of regions of interests

To determine ROIs, the Allen Mouse Brain Atlas (Allen Institute for Brain Science, http://mouse.brain-map.org/) (Lein et al., 2007) was spatially normalized with a mouse brain EPI template image using both affine and nonlinear transformations. All the ROIs that responded to visual stimuli were selected from the fMRI maps and the brain atlas: these were the LGd, SCs, LP, V1, V2, VISpor, and SUBcom, which correspond to the dorsal lateral geniculate nucleus, the sensory areas of the superior colliculus, the lateral posterior nucleus of thalamus, the primary visual area, the higher-order visual area, the postrhinal area, and the subiculum complex, respectively. The BOLD time courses of those ROIs were extracted from the fMRI data for each mouse and then averaged across all mice. The BOLD fMRI responses over the stimulation period were averaged in each animal, excluding the initial 3 s of data after stimulus onset to avoid the initial transition period.

#### Characteristics of the hemodynamic responses

To compare the dynamics of hemodynamic response functions (HRF) in the awake and anesthetized conditions (*Experiment #3*), a two-gamma-variate function often adopted for HRF estimation (Yu, Qian, Chen, Dodd, & Koretsky, 2014) was fitted to 40 data points right after the onset of the first stimulation in each ROI and each animal. Then, the times to 10% and 90% of the peak were calculated for each individual dataset and considered as the onset time and time to peak.

#### Frequency tuning

Frequency tuning curves were generated for each ROI from *Experiment #2*. At each ROI in each animal, the BOLD fMRI responses of all frequencies were normalized by the peak fMRI response to minimize inter-animal variations. Then, the temporal frequency tuning curve was obtained by plotting the normalized response versus the temporal frequency for each mouse, fitted with a Gaussian fitting function using Matlab®, and averaged across all mice. The preferred frequency was defined as the frequency of the peak response, and the bandwidth was obtained from the FWHM of the fitted tuning curve.

### 2.7. Statistics

All quantitative values are represented as the mean ± standard error of the mean (SEM). One-way repeated-measures analysis of variance (repeated ANOVA) followed by a Bonferroni post-hoc analysis was used for pulse duration and frequency-dependent BOLD fMRI, and independent t-testing was used to compare the BOLD responses in the awake vs. ketamine/xylazine conditions. The statistical significance of the results was assessed according to a p < 0.05 criterion.

## 3. Results

### 3.1. Movement of awake and anesthetized animals during fMRI

Head motion was assessed using six motion parameters (3 translation displacements along the x, y, and z axes and 3 rotational displacements of roll, pitch, and yaw) and FD (see Supplementary Fig.1 for a successful and failed trial in the awake condition). In the seven mice used for the awake fMRI, 66 of the 95 trials passed the FD threshold of 0.15 mm. We acquired 30 successful trials from 52 trials of the four mice from Group 1 (entirely trained in the mock scanner) and 36 successful trials from 43 trials of the three mice in Group 2 (partially trained in the actual MRI scanner). The data for one mouse in Group 1 were removed based on the FD criterion, and of the six remaining animals, 82.5% of the scans were successful (66/80 total scans). Only successful trials were included for further analysis.

The average time courses of the FD were plotted for six individual animals (Figure 2). The observed motions were not synchronized with the visual stimulation. Among the six awake animals, three were trained outside the scanner (Fig. 2A), and the remaining three were habituated inside the 9.4 T scanner for the last three days of training (Fig. 2B). A substantial reduction in head motions was observed among the animals acclimated to the actual MRI scanner (mean ± SEM average FD: 0.017 ± 0.0025 mm for inside the MRI scanner training vs. 0.04 ± 0.013 mm for outside magnet training).

**Figure 2.**
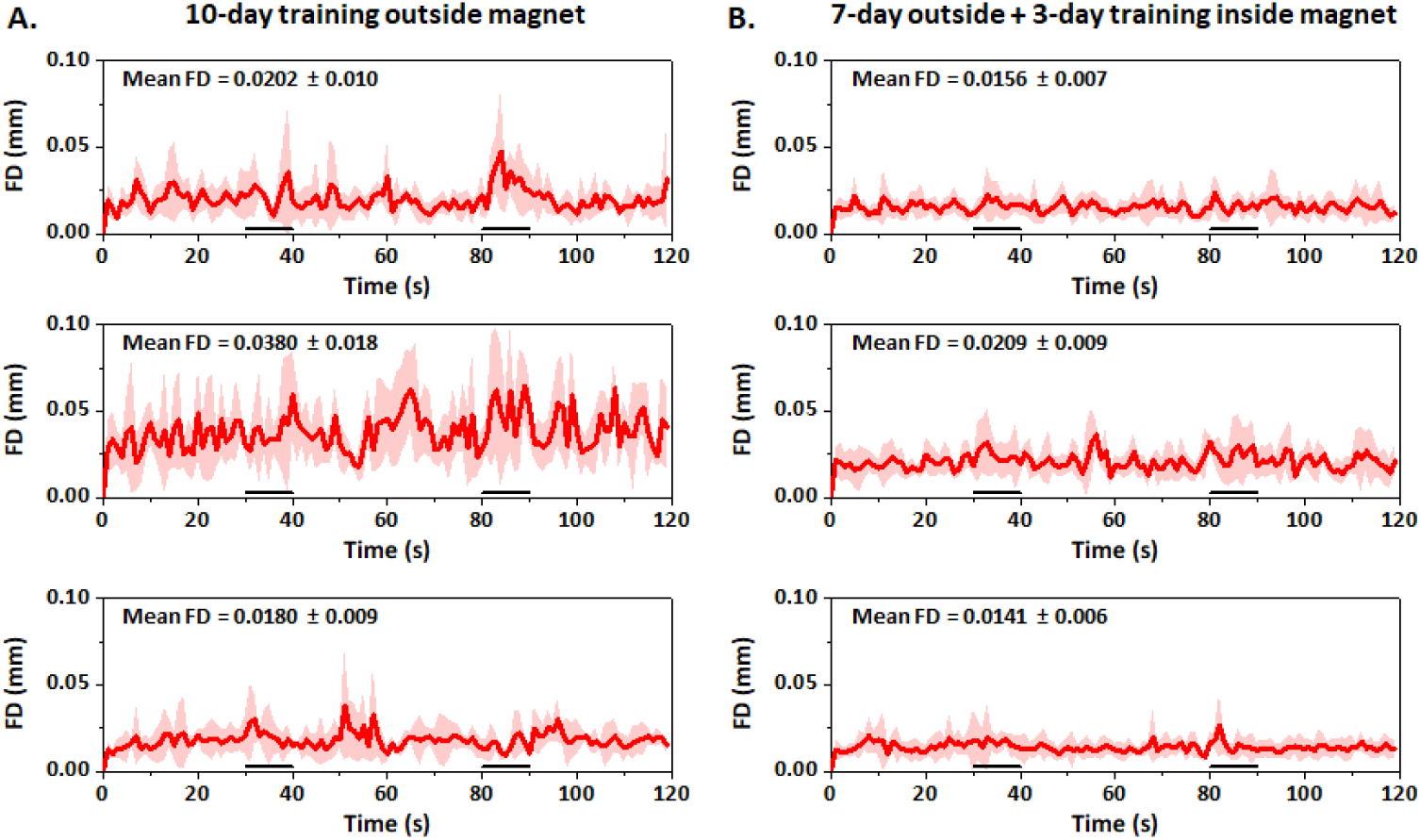
Animal-wise averaged time course of frame-wise displacements in awake mice trained using two different habituation procedures. All trials that passed the FD threshold of 1.5 mm were averaged in each animal. The three animals on the left side (A) underwent the total 10-day habituation outside the magnet (Group 1), whereas those on the right side (B) underwent the same habituation for 7 days plus 3 days in the 9.4 T MRI scanner (Group 2). The average of the mean FD values in each animal is reported in each panel. FD, frame-wise displacements; horizontal bars, stimulation periods; error bars, SEM.

The same estimation procedure was applied to the fMRI data from ketamine/xylazine anesthetized mice and the 3 Hz stimulation in *Experiment #2* to examine motion effects (Supplementary Fig. 2). The maximal translation was 0.004 ± 0.0004 mm along the y-axis (phase-encoding), and the maximal FD was 0.008 ± 0.001 mm, corresponding to 5.1% of the voxel dimensions (0.156 mm) and not synchronized to the visual stimulation. These small displacements were not noticeable. The respiratory rate ranged 160 – 180 beats per minute (bpm) without any noteworthy changes caused by the visual stimulation. However, to ensure consistency throughout all the data analyses, the motion regression was also applied to the anesthetized data.

### 3.2. Characteristics of BOLD responses with varying pulse length and stimulus frequencies under ketamine/xylazine anesthesia

To optimize the visual stimulation parameters and characterize the spatial patterns of BOLD responses, BOLD fMRI data were obtained during the presentation of a binocular flashing light stimulus under anesthesia.

The pulse duration–dependent responses to light stimulation were initially measured in seven mice at a fixed 3 Hz frequency. Significant bilateral positive BOLD responses were found in the main structures of the visual pathways (LGd, SCs, LP, V1, and V2) for all tested durations (Fig. 3A and Supplementary Fig. 3 for all frequencies), and they had similar patterns with slight variations. The BOLD response (Fig. 3B) was detectable within 2 – 3 s after the onset of stimulation, peaked at 5 – 8 s, and remained stable during the remaining stimulation period. After the cessation of stimulation, the BOLD signal decreased slowly over ∼20 s, and post-stimulus undershoot was observed in many conditions. The greatest response was observed in the SCs, which directly receives most retinal ganglion activity.

**Figure 3.**
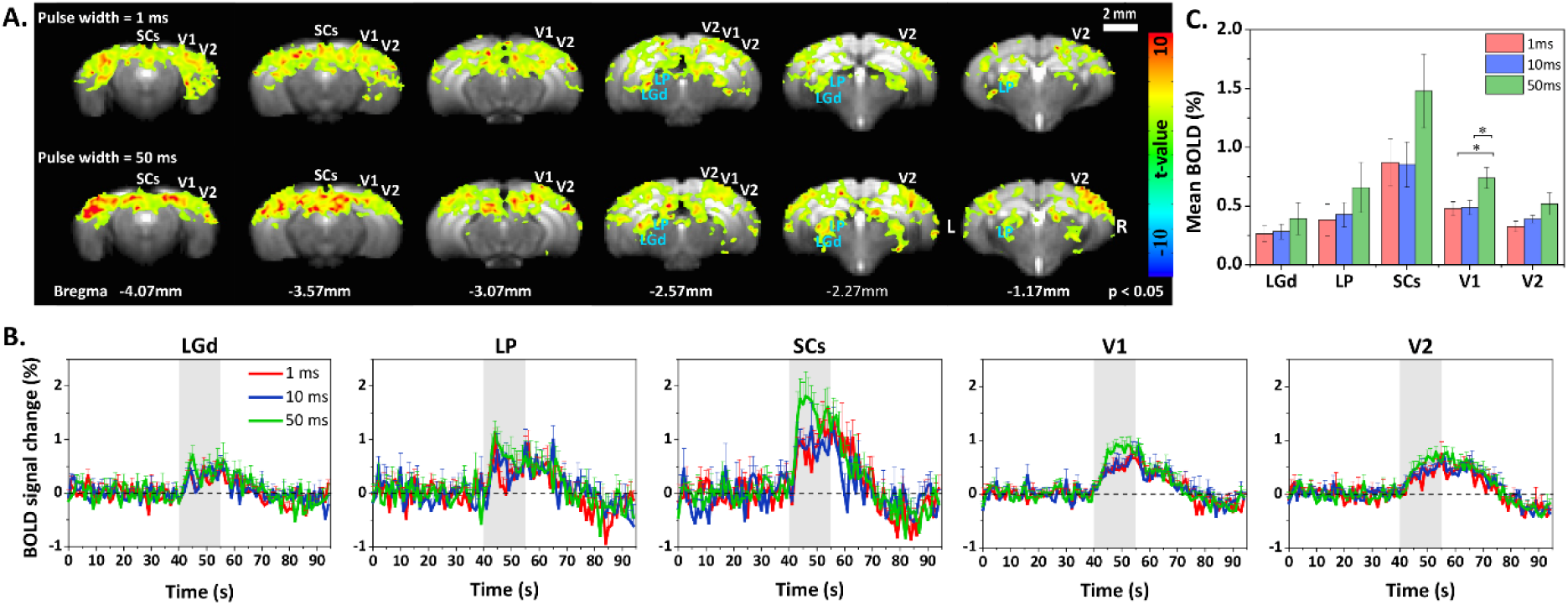
Pulse duration-dependent BOLD response to binocular visual stimulation under anesthesia. (A) Group activation maps of the visual stimulus with 1 ms and 50 ms pulse durations overlaid on mouse brain EPI template images (n = 7; p < 0.05, FWE corrected). Visual areas are labeled for better visualization, and the coronal slice positions are marked relative to Bregma. All pulse duration-dependent maps can be found in Supplementary Fig. 3. (B) Average time courses of visual ROIs for three different pulse durations. Error bars: SEM (n = 7); gray vertical bar: 15 s stimulation period. (C) Mean percent BOLD signal change in the LGd, SCs, LP, V1, and V2 for different pulse durations. Signal changes in V1 increased significantly with longer stimulation pulses (*p < 0.05; repeated measures ANOVA followed by Bonferroni post-hoc test).

The group results revealed that although increasing the pulse duration from 1 to 50 ms elicited a slight increase in the mean change in the BOLD signal, it was not statistically significant except in V1 (Fig. 3C). Overall, our results suggest that BOLD responses are not significantly influenced by varying the pulse duration from 1 to 50 ms.

Figure 4A illustrates the BOLD fMRI group activation maps from 3 Hz and 10 Hz stimulation under ketamine/xylazine anesthesia (n = 8 mice; Supplementary Fig. 4 contains all frequencies). During the 3 Hz stimulus, positive responses were robustly observed in the main ROIs of the visual pathway (LGd, SCs, LP, V1, and V2). During the 10 Hz stimulation, activation of the V1 and V2 cortical areas almost disappeared, and activation of the LGd, SCs, and LP areas was maintained but less pronounced than at the lower frequencies (Supplementary Fig. 4). The average time courses of BOLD responses corresponding to light stimuli at varying frequencies are shown in Fig. 4B (n = 8 mice). The BOLD fMRI response of the LGd to 10 Hz stimulation was the lowest, whereas the other frequencies all exhibited similar amplitudes. In the SCs, the amplitude of the BOLD responses to 1 Hz, 3 Hz, and 5 Hz stimuli was greater than that for 8 Hz and 10Hz stimuli (Fig. 4C). In the cortical areas, V1 and V2, the response increased as the stimulus frequency increased from 1 to 5 Hz; the response amplitude then decreased as the frequency was further increased to 8 and 10 Hz. When comparing the five areas assessed, the SCs had the most pronounced BOLD responses at all frequencies.

**Figure 4.**
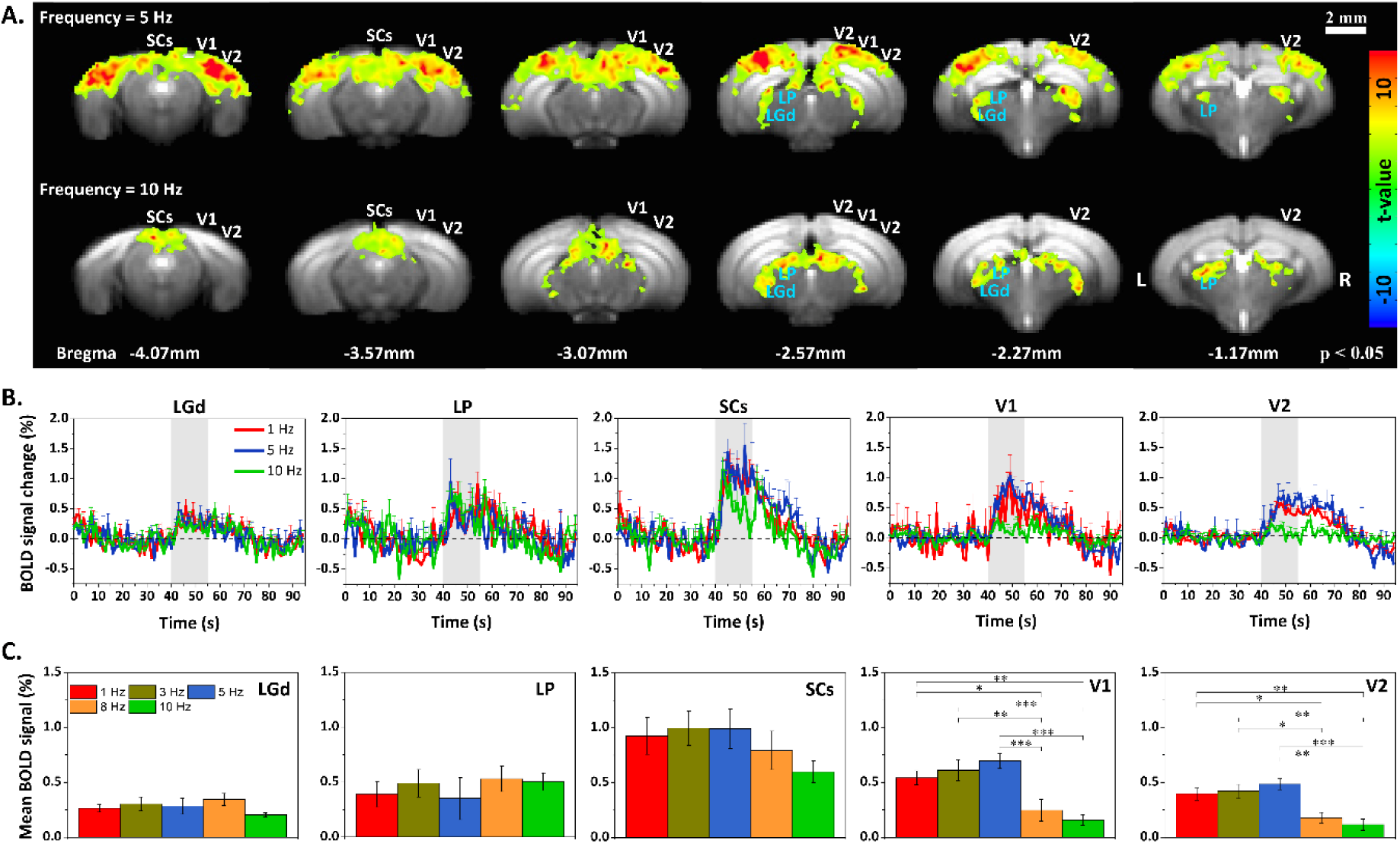
Frequency-dependent BOLD responses to binocular visual stimulation under anesthesia. (A) Group activation maps for the 5 and 10 Hz frequency stimuli (n = 8; p < 0.05, FWE corrected) overlaid on mouse brain EPI template images. Visual areas are labeled for better visualization, and coronal slice positions relative to the Bregma are marked. All pulse frequency–dependent maps can be found in Supplementary Fig. 4. (B) Averaged time courses and (C) mean percent changes of the BOLD responses in the visual ROIs at five stimulation frequencies. Error bars: SEM (n = 8); gray vertical bar: 15 s stimulation period; * p < 0.05, ** p < 0.01, *** p < 0.001 (repeated ANOVA followed by a Bonferroni post-hoc analysis).

### 3.3. fMRI responses of anesthetized vs. awake mice

Frequency-dependent BOLD responses were observed under ketamine and xylazine anesthesia. One important question is whether similar BOLD responses are detected under wakefulness. Based on the anesthetized fMRI studies, two stimulation frequencies (5 and 10 Hz) were selected for the comparison of fMRI data in the awake and anesthetized conditions. Figure 5 shows the fMRI group activation maps from 5 Hz stimulation in anesthetized mice (Fig. 5A), 5 Hz and 10 Hz stimulation in wakeful mice (Fig. 5B – C), and the time courses of both frequencies in both the awake and anesthetized conditions (Fig. 5D – E). Figure 6A shows the peak intensity response to 5 Hz stimulation under anesthesia and wakefulness. Observations are: 1) anesthesia induced diffuse BOLD responses compared with wakefulness; 2) BOLD responses in the SCs were similar in the awake and anesthetized conditions; 3) responses in the LGd were slightly stronger under wakefulness than under anesthesia; and 4) cortical responses in V1 and V2 were weaker under wakefulness than under anesthesia (Fig. 5D). When the stimulus frequency increased to 10 Hz, the BOLD response in V1 became negative (Fig. 5C – E).

**Figure 5.**
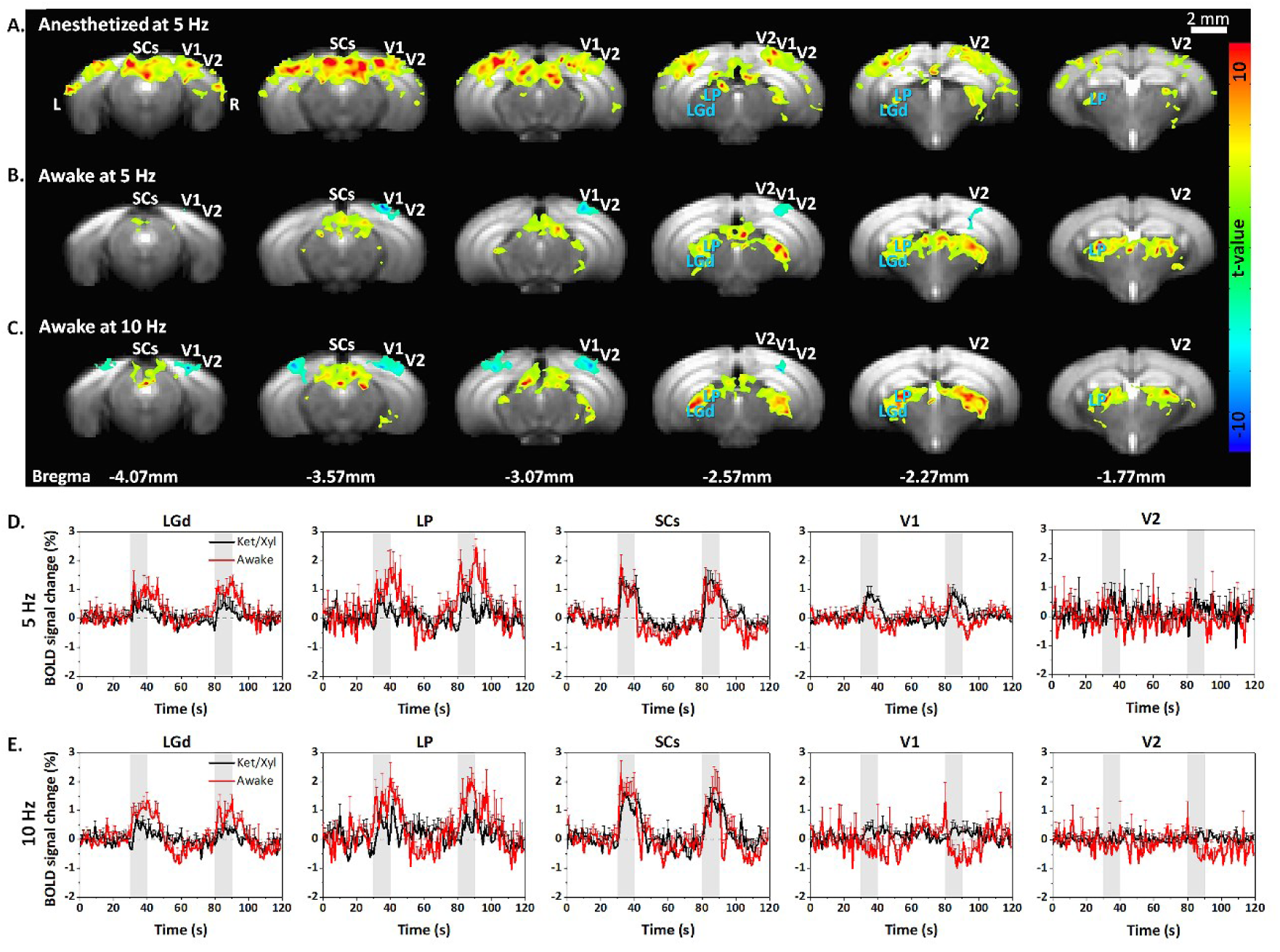
Functional activation maps and BOLD time courses following visual stimulation under the anesthetized and awake conditions. (A – B) Group activation maps following 5 Hz stimulation under anesthesia and wakefulness and (C) 10 Hz stimulation of 6 awake mice (p < 0.05, FWE corrected) overlaid on mouse brain EPI template images. (D – E) Averaged BOLD time courses in all visual ROIs in awake (red) and anesthetized (black) mice with 5 Hz (D) and 10 Hz (E) stimulation. Error bar: SEM; gray bars: 10 s visual stimulation.

**Figure 6.**
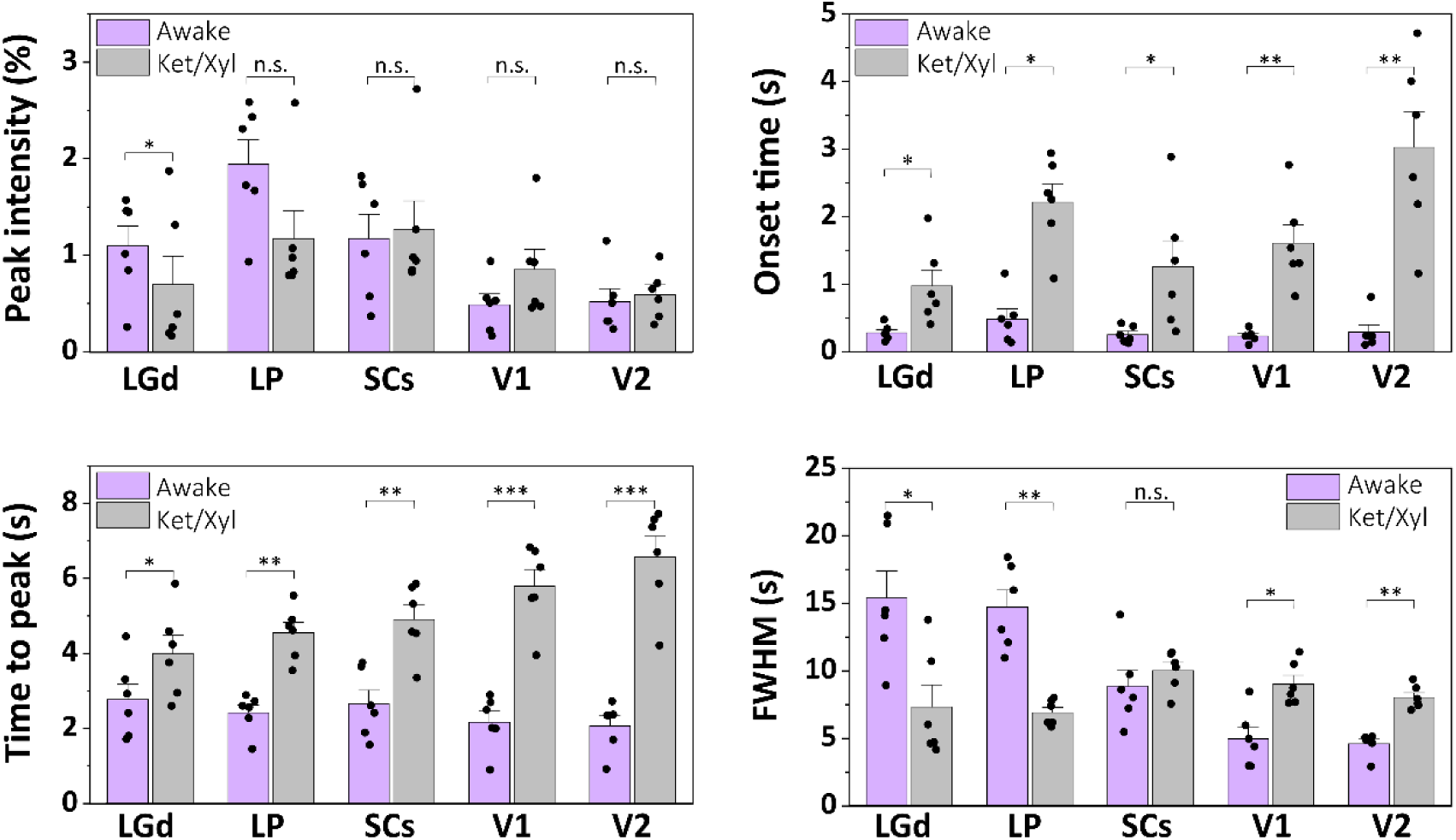
Characteristics of BOLD responses to 5 Hz visual stimulation under wakefulness and anesthesia. (A) The peak BOLD signal change at 5 Hz in the visual area. (B) Comparison of onset time, time to peak, and the full width at half maximum (FWHM) of BOLD responses under awake vs. ketamine/xylazine anesthesia conditions in *Experiment #3* (* p < 0.05, ** p < 0.01, *** p < 0.001).

#### Dynamics of BOLD responses between awake and anesthetized conditions

Times to 10% and 90% of the peak were calculated for each individual 5 Hz stimulation dataset and considered as the onset time and time to peak, respectively. BOLD responses during wakeful stimulation were 2–3 s faster than those from anesthetized mice in all regions (Fig. 6). The FWHM of the awake BOLD responses in the subcortical areas was significantly larger than that from anesthetized mice, but the cortical ROIs behaved oppositely.

### 3.4. Frequency tuning of mouse visual system

Frequency tuning is an important parameter in cellular properties. The normalized mean BOLD changes against stimulation frequencies and fitted frequency tuning curves in *Experiment #2* were calculated for five ROIs and are shown in Fig. 7A. The peak frequencies of the tuning curve under anesthesia were 4.3 ± 0.5 Hz and 4.4 ± 0.2 Hz for V1 and V2, respectively, 6.1 ± 0.8 Hz for the SCs, 6.8 ± 0. 7 Hz for the LP, and 6.2 ± 0.5 Hz for the LGd. The bandwidth of the tuning curve was 6.2 Hz for V1, 6.5 Hz for V2, 10.1 Hz for the SCs, 8.9 Hz for the LP, and 8.8 Hz for the LGd. The cortical ROIs thus had a lower preferred frequency and narrower tuning bandwidth than the subcortical ROIs (4.3 – 4.4 Hz vs. 6.1 – 6.8 Hz and 6.2 – 6.5 Hz vs. 8.8 – 10.1 Hz, respectively).

**Figure 7.**
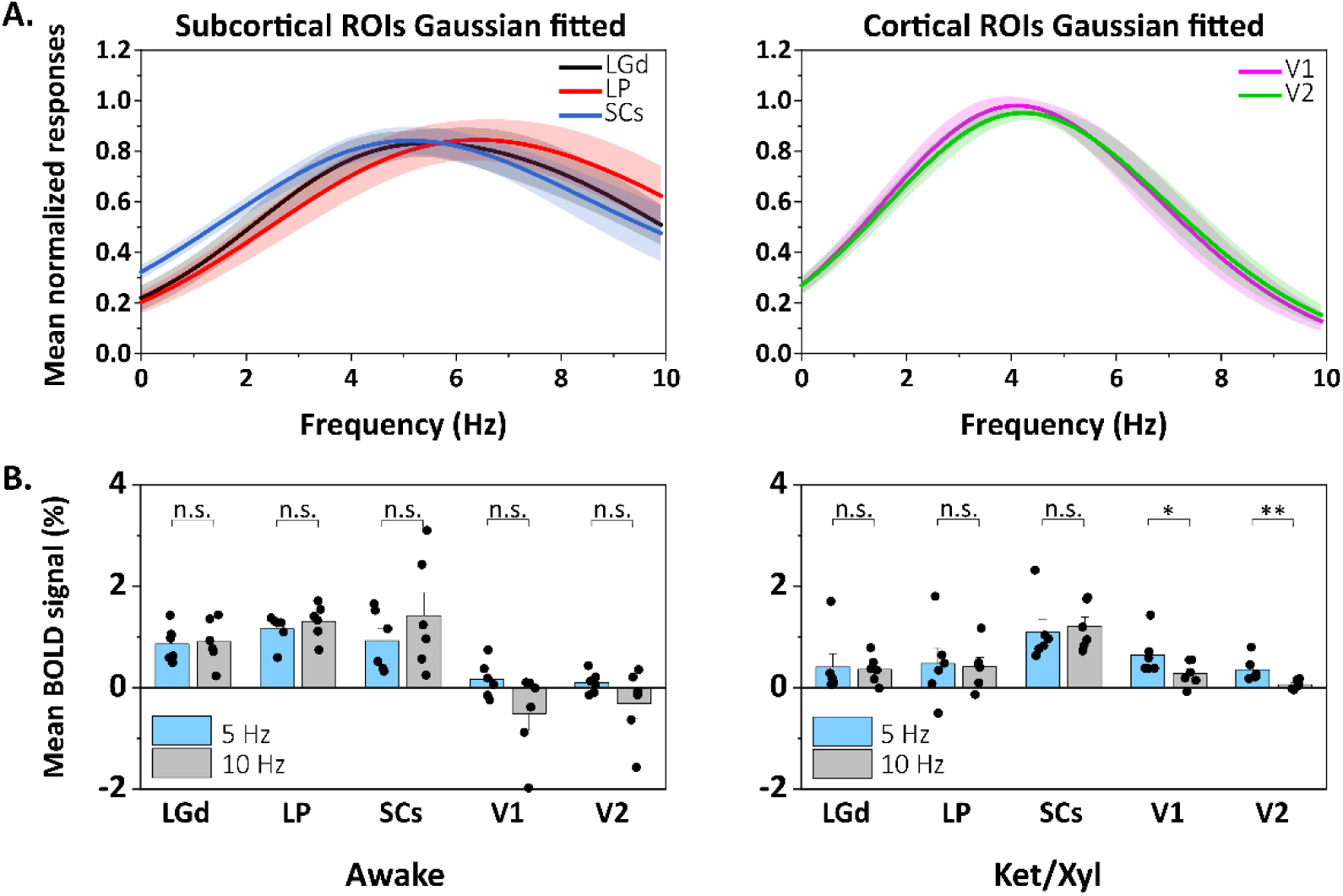
Frequency tuning properties of visual ROIs. (A) Normalized frequency-dependent response curves were obtained for subcortical ROIs and cortical ROIs in *Experiment #2*. The V1 and V2 responses overlapped completely. (B) Mean BOLD signals from awake vs. anesthetized mice in *Experiment #3*. Error bars: SEM (n = 8); * p < 0.05, ** p < 0.01, *** p < 0.001, n.s.: not significant.

To compare the visual responses from awake and anesthetized mice, mean BOLD signal changes at 5 Hz and 10 Hz were plotted in Fig. 7B. In the awake condition, the peak mean BOLD responses shifted toward a slightly higher frequency in the subcortical areas, with increased activity at 10 Hz compared with 5 Hz (the ratio of mean BOLD signals at 10 Hz to 5 Hz was 0.89 under anesthesia vs. 1.05 in wakefulness for LGd, 0.88 vs. 1.10 for LP, and 1.10 vs. 1.54 for SCs), but the cortical areas, V1 and V2, were fully suppressed. In general, the relative BOLD intensities under 5 Hz and 10 Hz stimulation were similar under wakefulness and anesthesia, indicating that the frequency tuning curve in both conditions is similar.

### 3.5. Activation in non-visual areas

In addition to the bilateral BOLD fMRI responses in the main visual pathway areas, distinct BOLD fMRI responses to visual stimulation were observed in unexpected regions, the postrhinal area (VISpor) and the subiculum complex (SUBcom, which includes the pre-subiculum and post-subiculum), under ketamine/xylazine anesthesia. Figure 8 shows the activation maps responding to 5 Hz stimulation under ketamine and xylazine and the frequency tuning curves of those two areas. Under ketamine/xylazine anesthesia, the frequency tuning of the VISpor and SUBcom (Fig. 8B) appeared similar to that of V1 and V2, with the peak activation at 4.8 Hz and 3.7 Hz and a bandwidth of 6.9 Hz and 4.1 Hz, respectively. However, in the awake condition (Fig. 8C), the BOLD response in these two ROIs disappeared, indicating that the anesthesia evoked the BOLD activity in these areas.

**Figure 8.**
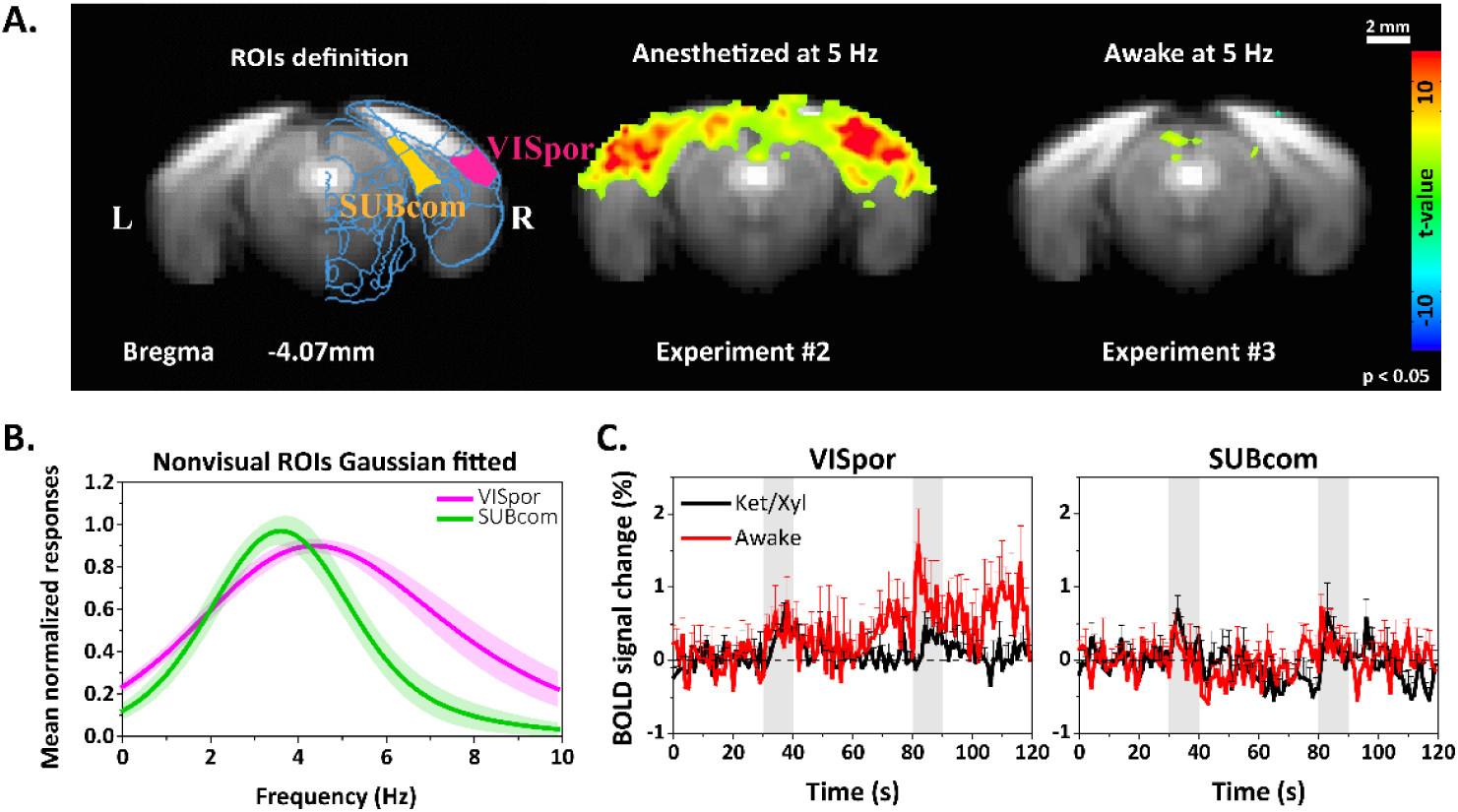
Activation in the postrhinal area and subiculum complex during visual stimulation under anesthesia. (A) ROI definitions overlaid on an EPI template of the postrhinal area and the non-visual subiculum complex (left) and BOLD activation maps following 5 Hz visual stimulation under ketamine/xylazine anesthesia (middle) and awake (right) (p < 0.05, FWE corrected). (B) Frequency tuning curves of non-visual areas under ketamine/xylazine from *Experiment #2*. (C) Time courses of the postrhinal area and subiculum complex responding to 5 Hz stimulus from the *Experiment #3* comparison between anesthetized and wakeful mice. In awake animals, the fMRI responses in these two non-visual areas were reduced. Error bars: SEM; vertical gray bars: 10 s visual stimulation.

## 4. Discussion

We investigated the characteristics of BOLD functional activation in response to visual stimulation with light of different pulse durations and frequencies in ketamine and xylazine anesthetized mice. We also developed an awake mouse protocol and compared the BOLD fMRI responses to visual stimulation under wakefulness and ketamine/xylazine anesthesia. Our main findings are as follows. (1) BOLD fMRI activities occurred in all visual pathway-related regions under both awake and anesthetized conditions. In all regions, the BOLD response was faster under wakefulness than anesthesia, as expected. The subcortical areas (LGd, LP, SCs) had higher BOLD responses under wakefulness than in the anesthesia condition, whereas the cortical areas (V1 and V2) produced biphasic/negative BOLD responses in the awake condition and positive BOLD responses under ketamine/xylazine anesthesia. (2) In terms of visual stimulation parameter modulations, the effect of pulse duration ranging between 1 and 50 ms was not significant. However, stimulation frequency did significantly affect BOLD responses. The mouse visual system is sensitive to low-frequency visual stimulation (∼ 4 Hz). The subcortical areas responded to higher stimulation frequency with broader FWHM compared with the cortical areas. (3) We also observed activation in the extrastriate visual postrhinal area and the subiculum complex, including the post-subiculum and pre-subiculum, under anesthesia that almost disappeared under wakefulness.

### 4.1. fMRI response in awake vs. anesthetized conditions

We have successfully developed an awake fMRI protocol by designing a restraint apparatus and habituation protocol. Many attempts have been made to perform awake mouse fMRI to minimize the effects of anesthesia on neural activity and neurovascular coupling (Chen et al., 2020; Desai et al., 2011; Desjardins et al., 2019; Harris et al., 2015; Schlegel et al., 2015; Yoshida et al., 2016). The fundamental requirement for studying awake animals is a restraint apparatus that can minimize head motion and reduce artifacts associated with body motion. Minimization of head motion can easily be achieved by adopting the head post, but the procedure to attach the head post to the skull often induces image artifacts through residual blood clots and trapped air bubbles. To minimize the translation of body motions to the head, most published studies tried to use a long tube fitted to the animal body, and some tried to secure all animal limbs using surgical tape, with the animal’s incisors secured over a bite bar (Chen et al., 2020; Desai et al., 2011; Desjardins et al., 2019; Han et al., 2019; Harris et al., 2015; Tsurugizawa et al., 2020; Yoshida et al., 2016). All those procedures could induce more stress and artifacts from body movement. We designed a detachable cover piece to tightly fit to the neck (Fig. 1A). The major innovation of our cradle is open space for the four paws (Fig. 1A), which allows the paws to move freely without inducing significant body motions and allows our cradle to be used for detecting paw responses to behavioral tasks.

In our study, we used a 10-day acclimation protocol to reduce animal stress levels and motion. In our preliminary studies (data not shown), we measured corticosterone levels (as a stress marker) in the blood of two mice and found them to be ∼10ng/ml before training started, 30ng/ml on the 5th day, and 15ng/ml after the 10th day of outside-MRI habituation, suggesting that the stress level was reduced by habituation but remained modest. Even under acclimation, animal restraints can cause a higher stress level than the normal condition in mice (Gong et al., 2015; Tsurugizawa et al., 2020). Moving the habituation procedure to the actual MRI scanner for last 3 days reduced movements and increased the success rate of the awake fMRI scans. Even though the magnet training procedure required expensive imaging time and resources, it was useful to acclimate the animals to the fMRI environment because it increased the rate of successful fMRI scans.

In both the awake and anesthetized conditions, BOLD fMRI activity was observed in all visual pathway related areas. All the BOLD responses of awake animals were faster than those under ketamine and xylazine anesthesia, which is consistent with observations by Desai et al. (Desai et al., 2011). Based on previous awake vs. anesthetized fMRI studies, we also expected to have reduced BOLD responses under anesthesia; in earlier work BOLD responses to somatosensory stimulation were reduced about 3 times under 0.7% isoflurane compared with awake responses (Desai et al., 2011). However, under ketamine and xylazine anesthesia, the BOLD signal was reduced only slightly in the subcortical regions (Fig. 6A). The most interesting observation was in the V1 activity. Under ketamine/xylazine anesthesia, the BOLD activity in V1 was sustained during the stimulation period and spread into a large area beyond V1, whereas the awake BOLD response was biphasic or negative (Fig. 5). A biphasic response to 5 Hz stimulation in V1 under the awake condition produced negative BOLD maps when a standard hemodynamic response function was used to generate the fMRI maps (Fig. 5B, V1 area). Our BOLD fMRI observation can be explained by previous investigations of visually evoked synaptic responses in the V1 of urethane-anesthetized vs. awake mice (Haider et al., 2013; Vaiceliunaite et al., 2013). Under anesthesia, synaptic responses occurred in a large region of the visual area and were prolonged by the reduced inhibition of neural activity, whereas under wakefulness, visually evoked responses were more spatially selective and much briefer due to stronger inhibition than excitation. Thus, our BOLD data are consistent with previous neurophysiological data (Haider et al., 2013; Vaiceliunaite et al., 2013).

The mixture of ketamine and xylazine keeps animals in a near-arousal state (Shim et al., 2018). Ketamine tends to bind selectively to the NMDA (N-methyl-d-aspartate receptor antagonist) receptors of GABAergic inhibitory interneurons (Olney, Newcomer, & Farber, 1999; Seamans, 2008), resulting in the disinhibition of pyramidal cells (Fan et al., 2018). Therefore, the suppression of inhibitory activities and sensitization of excitatory activities could increase the functional sensitivity of neurons, especially those in the cortical regions where inhibitory and excitatory neural connections are dense. The sensitization of excitatory neurons by ketamine sustains and spatially diffuses the cortical BOLD response.

Our observations have important implications for fMRI mapping. 1) A biphasic or negative response in cortical regions with a delicate excitation and inhibition balance can be detected due to stronger inhibition than excitation. An initial positive BOLD response is likely induced by initial excitation, whereas a negative BOLD signal is caused by delayed inhibition, as seen in synaptic measurements (Haider et al., 2013; Vaiceliunaite et al., 2013). If the anesthetic used for fMRI is a GABAergic receptor agonist such as isoflurane, cortical inhibition is enhanced, resulting in negative or no responses to evoked neural stimulation. Negative or no V1 activity in response to visual stimulation in our awake studies and medetomidine-anesthetized fMRI (Niranjan et al., 2016) can be explained by larger inhibition-than-excitation effects. 2) Prolonged and diffused V1 activity under ketamine/xylazine anesthesia possibly projects to other regions including the V2, postrhinal area, and subiculum complex. The advantage of using ketamine is enhancing excitatory activities by disinhibition, which produces increased BOLD responses in the primary sensory cortex and its projected regions. Consequently, widely distributed functional brain networks can be mapped under ketamine/xylazine, indicating that the mixture of ketamine and xylazine is a good anesthetic for mapping functional networks in mouse fMRI.

### 4.2. Dependence of visual stimulation frequency under ketamine/xylazine anesthesia

The neurons of the mouse visual system respond selectively to specific types of visual stimuli (Hubel & Wiesel, 1962; Hubener, 2003; Morin & Studholme, 2014). Thus, temporal frequency is a fundamental attribute of visual stimuli that relates directly to visual sensitivity, and studying the regional temporal frequency tuning properties of the mouse visual system is a first step toward understanding the processing of visual information.

The tuning properties measured by fMRI under ketamine/xylazine anesthesia were consistent with those previously reported in electrophysiology studies of mice (Durand et al., 2016; Grubb & Thompson, 2003; Piscopo et al., 2013). In the LGd, the fMRI peak frequency under anesthesia was 6.2 Hz with an FWHM of 8.8 Hz, which was similar to or slightly higher than that in published electrophysiology studies, in which the tuning frequency was 6.3 Hz for wakefulness, 4.1 Hz for urethane anesthesia (Durand et al., 2016), and ∼4 Hz with an FWHM of 7 Hz under halothane and nitrous oxide anesthesia (Grubb & Thompson, 2003). The LP tuning was similar to that of the LGd, with a tuning frequency of 6.8 Hz and an FWHM of 10.1 Hz, which concurred with previously published electrophysiology results from awake mice (6.3 ± 0.2 Hz), in which the LP was also found to have values similar to those of the LGd (Durand et al., 2016). Not only the LP but SCs neurons produce various responses to different visual stimuli; however, the main characteristics of the responses are also similar to those of the LGd (Gale & Murphy, 2014). These properties can explain our results, in which the SCs shared an analogous tuning frequency with the LGd, with a high preferred frequency of 6.1 Hz. In the cortical areas (V1 and V2), the peak tuning frequency of 4.3 – 4.4 Hz was slightly higher than in the published electrophysiology studies, in which a 3 Hz tuning frequency at V1 was obtained in both awake and urethane anesthetized animals (Durand et al., 2016), and a tuning frequency of ∼4 Hz was found at layers 2/3 and 4 under urethane anesthesia (Niell & Stryker, 2008). Because V1 activity is transferred to the higher-order V2 (Felleman & Van Essen, 1991; Kalatsky & Stryker, 2003; Wagor, Mangini, & Pearlman, 1980; Q. Wang & Burkhalter, 2007), it was expected that the properties of the V1 and V2 would be similar. Because the bandwidth for the frequency tuning in V1 and V2 was 6.2 – 6.5 Hz, the fMRI response of the cortical areas was strongly suppressed at high frequencies (8 Hz and 10 Hz). The slight differences between our results and the electrophysiology data could have resulted from multiple issues, including the resolution of temporal frequencies and the sensitivity and signal sources of the techniques used. The tuning frequency in ketamine/xylazine anesthetized mice is consistent with that obtained by BOLD fMRI in isoflurane-anesthetized cats, 3.1 – 4.5 Hz for V1 and V2 and 6 Hz for LGd (Yen, Fukuda, & Kim, 2011). In sum, our frequency tuning results under ketamine/xylazine anesthesia agree well with the electrophysiology data, indicating that the mixture of ketamine and xylazine is a good anesthetic for mouse fMRI.

### 4.3. Comparison with previous rodent fMRI studies

Only a few previous mouse fMRI studies have used visual stimulation, and their outcomes differ (Huang et al., 1996; Lee et al., 2019; Niranjan et al., 2016). In the Huang at al. mouse fMRI study (Huang et al., 1996), photic stimulation of sodium pentobarbital anesthetized mice produced positive responses encompassing V1 and V2, whereas the superior neighbor displayed a negative response upon stimulation (Huang et al., 1996). However, due to the limited spatial resolution achieved in that study (0.23 × 0.46 × 2 mm^3^), clear demarcation of different visual regions was not possible. Niranjan et al. (Niranjan et al., 2016) reported mouse fMRI data with a spatial resolution of 0.36 × 0.36 × 0.5 mm^3^ obtained under medetomidine sedation and found positive BOLD responses in the V1, LGd, and SCs. At 5 – 10 Hz, they obtained negative BOLD responses in V1 but not outside the primary visual areas. Recently, Lee et al. (Lee et al., 2019) measured BOLD fMRI responses to 5 Hz visual stimulation in medetomidine/isoflurane anesthetized mice and found a peak intensity of 0.3% in V1 (TE = 15 ms) (Fig. 4 in Lee, Li, Coulson, & Chuang, 2019), which is less than our observation (Fig. 5A) under ketamine/xylazine anesthesia.

The frequency dependence of activation in the mouse visual areas that we found concurs with that found in rat studies (Bailey et al., 2013; Pawela et al., 2008; Van Camp, Verhoye, De Zeeuw, & Van der Linden, 2006), in which the subcortical areas had positive BOLD responses, and the cortical areas tended to have reduced responses with increasing stimulus frequency. Those rat data indicate that subcortical areas have broad high-frequency tuning, whereas the cortical areas have narrow low-frequency tuning. This tendency somewhat agrees with previously published electrophysiology data and our fMRI data, but the specifics of such patterns can be quite different. For example, Bailey et al. performed BOLD fMRI in urethane-anesthetized rats responding to 1, 5, and 10 Hz LED stimulation and found increasing BOLD responses in the SCs with increasing stimulation frequency (Bailey et al., 2013). In our study, the BOLD responses in the subcortical areas were reduced at high frequencies (8 Hz and 10 Hz), but not as much as in the cortical areas. All those discrepancies can be explained by the species differences between mice and rats and the anesthetic used (our results vs. those of Niranjan et al., 2016).

### 4.4. Observation of BOLD fMRI responses in non-visual areas

In addition to detecting activation in the main visual areas, we also detected activation in the retinotopically organized postrhinal area (VISpor), which is known as one of the ventral extrastriate visual areas in mice. The VISpor receives visual information from V1 (Beltramo & Scanziani, 2019; Glickfeld & Olsen, 2017; Nishio et al., 2018; Q. Wang & Burkhalter, 2007), and is involved in recognizing visual objects and encoding spatial information in the environment (Furtak, Ahmed, & Burwell, 2012). Also, the VISpor responds to static stimuli and is robustly modulated by moving stimuli (Nishio et al., 2018). The BOLD response of the VISpor was similar to that of V1. Under ketamine/xylazine anesthesia, the BOLD response of the VISpor was strong upon low-frequency light stimulation and suppressed at high frequencies. In the awake condition, the BOLD response of the VISpor was weak and decayed sharply after stimulation onset, resulting in no significant activation (Fig. 8).

Visual information can project from visual regions to the somatosensory, retrosplenia, cingulate, orbitofrontal, temporal, and parahippocampal cortexes (Q. X. Wang, Sporns, & Burkhalter, 2012). But activation in those projected regions is not detected by fMRI. An unexpected finding in this study was activation in a non-visual area, the SUBcom, including the post-subiculum and pre-subiculum. Both the post-subiculum and pre-subiculum are known to play critical roles in navigation (Ding, 2013), and they are thought to connect to the visual cortex based on the findings of a previous electrophysiology study (Beltramo & Scanziani, 2019). Under anesthesia with a long stimulation duration, 15 s in *Experiments #1* and *#2*, the SUBcom was strongly activated and tuned to a frequency similar to that of V1 and V2, but the BOLD activity in the SUBcom disappeared in wakefulness, like that of the VISpor (Fig. 8). The visual cortex sends visual information to the hippocampus through the subiculum (Sun et al., 2019) for learning, but in our experiment only a simple flashing light was presented. The activation of the SUBcom was thus likely caused by the disinhibition property of the anesthetics we used. Consequently, visual information might be conveyed to functionally connected regions where evoked activities are not generally detected when neural activity is strongly inhibited.

### 4.5. Limitations and future studies

Our study had several limitations. First, we used simple LED stimulation, which is a rudimentary paradigm for exploring the mouse visual system. Thus, a better paradigm, such as moving gratings, is needed to improve fMRI investigations of visual networks. Second, we imaged a 4.5-mm coronal slab every second. This posterior-anterior coverage does not include the prefrontal cortex (i.e., the frontal eye field), which modulates visual processing (Zanto, Rubens, Thangavel, & Gazzaley, 2011). To further investigate visual processing, it is necessary to image the entire brain with a fast temporal resolution using advanced technology, such as simultaneous multi-slice EPI with a single coil (Lee et al., 2019). Third, to dissect visual networks, fMRI should be combined with molecular tools such as optogenetic/chemogenetic manipulations to silence or enhance activities in certain visual areas.

## 5. Conclusion

In this study, an awake fMRI procedure for mice was developed, and the mouse visual system was successfully mapped with BOLD fMRI under ketamine/xylazine anesthesia and wakefulness. Our BOLD fMRI observations are highly compatible with previously reported electrophysiological findings (Durand et al., 2016; Grubb & Thompson, 2003; Huberman & Niell, 2011; Piscopo et al., 2013), suggesting that ketamine/xylazine is an acceptable anesthetic for mouse fMRI, which has been shown to be a viable tool for investigating the relationships between genes and neurological functions in transgenic mice by determining cell-type-specific neural networks with multimodal fMRI and mapping functional reorganizations in health and disease.

## Acknowledgments

We would like to thank Dr. Joonyeol Lee and Mr. Andrew You for their contributions to our scientific discussions, Drs. Sangwoo Kim and Jungryun Lee for implementing initial awake fMRI protocol, Mr. Chanhee Lee for maintaining the MRI instruments, Ms. Sun Myung Park and Jung-mi Lee for their preparation of the animals for the awake fMRI studies.

